# Orthogonal neural representations support perceptual judgements of natural stimuli

**DOI:** 10.1101/2024.02.14.580134

**Authors:** Ramanujan Srinath, Amy M. Ni, Claire Marucci, Marlene R. Cohen, David H. Brainard

**Author notes:** Corresponding Author: David H. Brainard. equal contribution (Ramanujan Srinath, Amy M. Ni). equal contribution (Marlene R. Cohen, David H. Brainard).

## Abstract

In natural behavior, observers must separate relevant information from a barrage of irrelevant information. Many studies have investigated the neural underpinnings of this ability using artificial stimuli presented on simple backgrounds. Natural viewing, however, carries a set of challenges that are inaccessible using artificial stimuli, including neural responses to background objects that are task-irrelevant. An emerging body of evidence suggests that the visual abilities of humans and animals can be modeled through the linear decoding of task-relevant information from visual cortex. This idea suggests the hypothesis that irrelevant features of a natural scene should impair performance on a visual task only if their neural representations intrude on the linear readout of the task relevant feature, as would occur if the representations of task-relevant and irrelevant features are not orthogonal in the underlying neural population. We tested this hypothesis using human psychophysics and monkey neurophysiology, in response to parametrically variable naturalistic stimuli. We demonstrate that 1) the neural representation of one feature (the position of a central object) in visual area V4 is orthogonal to those of several background features, 2) the ability of human observers to precisely judge object position was largely unaffected by task-irrelevant variation in those background features, and 3) many features of the object and the background are orthogonally represented by V4 neural responses. Our observations are consistent with the hypothesis that orthogonal neural representations can support stable perception of objects and features despite the tremendous richness of natural visual scenes.

**Significance Statement:** We studied how the structure of the mid-level neural representation of multiple visual features supports robust perceptual decisions. We combined array recording with parametrically controlled naturalistic images to demonstrate that the representation of a central object’s position in monkey visual area V4 is orthogonal to that of several background features. In addition, we used human psychophysics with the same stimulus set to show that observers’ ability to judge a central object’s position is largely unaffected by variation in the same background features. This result supports the hypothesis that orthogonal neural representations can enable stable and robust perception in naturalistic visual environments and advances our understanding of how visual processing operates in the real world.

## Introduction

A major function of the visual system is to infer properties of currently relevant stimuli, without interference from the tremendous amount of task-irrelevant information that bombards our retinas. Many laboratory studies of the neural basis this ability use, for good reasons, simple (von der Heydt et al., 1984; Peterhans and Heydt, 1991; Gallant et al., 1993; Leopold and Logothetis, 1996; Pasupathy and Connor, 2002; Rust and Movshon, 2005; Martinez-Garcia et al., 2019; Peters and Kriegeskorte, 2021; Snow and Culham, 2021). An advantage of this approach is experimental control: one can parametrically vary stimuli and completely specify the input to the visual system. A downside of using such stimuli, however, is that their very simplicity prevents them from fully illuminating the neural algorithms by which the brain sorts through the large quantity of visual information that is characteristic of natural viewing (see simulations in (Ruff et al., 2018a)).

In contrast to simple artificial stimuli, natural images can vary in many features, and these features are jointly encoded by the responses of populations of neurons in visual cortex (Cadieu et al., 2007; Oleskiw et al., 2018; Kim et al., 2019; Yamane et al., 2020; Srinath et al., 2021; Hatanaka et al., 2022). To investigate the relationship between the representation of multiple features within such populations, we consider a high-dimensional neural-response space in which each dimension represents the firing rate of one neuron (Kohn et al., 2020; Vyas et al., 2020). The response of the entire population at any given moment (e.g. in response to one visual scene) is a point in this space. Systematically varying one scene feature, such as the position of a banana, traces out a continuous trajectory in the response space, which can typically be approximated by a line (Misaki et al., 2010; Okazawa et al., 2021a). If just one scene feature varies, then that feature can be read out by projecting the response onto this line (aka linear decoding). If multiple features can vary, they can each be linearly decoded without interference from the others if their variation traces out orthogonal lines. But robust readout will be difficult, if not impossible, if multiple features trace out similar lines. Intermediate cases are also possible, in which robust readout is possible but requires processing more complex than linear projection (e.g., quadratic classification (Burge, 2020)).

Following Hong et al., 2016, we reasoned that variation in task-irrelevant features of a natural scene should not impair performance on a visual task if two conditions are met: visual information is read out of a neural population in a way that approximates a linear decoder, and the representations of relevant and irrelevant features are orthogonal in the relevant neural populations.

Here, we leverage the power of computer graphics to take parametric control of stimulus features in naturalistic stimuli, enabling us to vary many naturalistic stimulus dimensions and test the hypothesis that the observers’ perceptual abilities to make fine perceptual distinctions will not be perturbed on a threshold-level judgment task by task-irrelevant variations in stimuli and backgrounds if the neural representations of task-relevant and irrelevant features are orthogonal. Using a combination of human psychophysics and monkey neurophysiology, we demonstrate that 1) the population representation of object position in V4 is orthogonal to those of several background features, 2) the ability of human subjects to make precise perceptual judgments about object position was largely unaffected by task-irrelevant variation in those background features, and 3) many features of the object and the background (position, color, luminance, rotation, and depth) are independently decodable from V4 population responses. Together, these observations support the idea that orthogonal neuronal representations enable stable perception of objects and features despite the tremendous irrelevant variation inherent in natural scenes.

## Materials and Methods

### Data Availability Statement

The data and code that generate the figures in this study have been deposited in a public Github repository https://github.com/ramanujansrinath/UntanglingBananas. MATLAB code for creating and displaying the images for human psychophysical experiments, as well as analyzing the raw data from these experiments, can be found at https://github.com/AmyMNi/NaturalImageThresholds. Request for further information should be directed to and will be fulfilled by the corresponding author David H. Brainard in consultation with the other authors.

### Experimental Models and Subject Details

#### Monkey electrophysiology

Two adult male rhesus monkeys (Macaca mulatta, 10 and 11 kg) were implanted with titanium head posts before behavioral training. Subsequently, multielectrode arrays were implanted in cortical area V4 identified by visualizing the sulci and using stereotactic coordinates. All animal procedures were approved by the Institutional Animal Care and Use Committees of the University of Pittsburgh and Carnegie Mellon University.

#### Human Psychophysics

This study was preregistered at ClinicalTrials.gov, NCT number NCT05004649, https://clinicaltrials.gov/ct2/show/NCT05004649. The experimental protocols were approved by the University of Pennsylvania Institutional Review Board. Participants were invited to volunteer to participate in this study. Participants provided informed consent and filled out a lab participant survey. We also screened for visual acuity using a Snellen eye chart and for color deficiencies using the Ishihara plate test. Participants were excluded prior to the experiment if their best-corrected visual acuity was worse than 20/40 in either eye or if they made any errors on the Ishihara plate test.

Participants were excluded after the conclusion of their first session if their horizontal position discrimination threshold in the no variation condition (see description of conditions below) was higher than 0.6 degrees of visual angle, and participants excluded at this point did not participate in any further experimental sessions.

### Experimental Design

#### Image Generation (for both human psychophysics and monkey electrophysiology)

All the stimuli were variants of the same natural visual scene: a square image with a central object (a banana) presented on an approximately circular array of overlapping background objects (made up of overlapping branches and leaves). The central object and/or the background objects changed in horizontal position, rotation, and/or depth across different stimuli. In the larger set of stimuli (detailed below) the luminance and color of the central and background objects also changed. The central object and background objects are presented in the context of other objects (a rock ledge, a skyline, and three moss-covered stumps) that remain unchanged across all stimulus conditions. This natural visual scene was created using Blender, an open-source 3D creation suite (https://www.blender.org, Version 2.81a). The object and background parameters were varied using ISET3d, an open-source software package (https://github.com/ISET/iset3d) that works with a modified version of PBRT (https://github.com/scienstanford/pbrt-v3-spectral; unmodified version at https://github.com/mmp/pbrt-v3).

The images created using ISET3d were converted to RGB images using custom software (Natural Image Thresholds; https://github.com/AmyMNi/NaturalImageThresholds) written using MATLAB (MathWorks; Natick, MA) and based on the software package Virtual World Color Constancy (github.com/BrainardLab/VirtualWorldColorConstancy). Natural Image Thresholds is dependent on routines from the Psychophysics Toolbox (http://psychtoolbox.org), ISET3d (https://github.com/ISET/iset3d), ISETBio (http://github.com/isetbio/isetbio), PBRT (https://github.com/scienstanford/pbrt-v3-spectral; unmodified version at https://github.com/mmp/pbrt-v3), and the Palemedes Toolbox (palamedestoolbox.org).

To convert a hyperspectral image created using ISET3d to an RGB image for presentation on the calibrated monitor, the hyperspectral image data were first used to compute LMS cone excitations. The LMS cone excitations were converted to a metameric rendered image in the RGB color space of the monitor, based on the monitor calibration data. A scale factor was applied to this image so that its maximum RGB value was 1 and the image was then gamma corrected, again using monitor calibration data. This process was completed separately for the two different monitors used, one for the psychophysics and one for the neurophysiology.

### Monkey electrophysiology

#### Array implantation, task parameters

Both animals were implanted with titanium headposts prior to behavioral training. After training, microelectrode arrays were implanted in area V4 (96 recording sites; Blackrock Microsystems). Array placement was guided by stereotactic coordinates and visual inspection of the sulci and gyri. The monkeys were trained to perform a fixation task along with other behavioral tasks that were not relevant to this study. The stimulus images used in this study were not displayed outside of the context of this task. The monkeys fixated a central spot for a pre-stimulus blank period of 150-400ms followed by stimulus presentations (200-250ms) interleaved with blank intervals (200-250ms). The stimuli were presented one at a time at a peripheral location that overlapped the receptive fields of the recorded neurons. In each trial, 6-8 stimuli were presented, after which the monkey received a liquid reward for having maintained fixation on the central spot until the end of the stimulus presentations. If the monkey broke fixation before the end of the stimulus presentations, the trial was terminated. The intertrial interval was at least 500ms. The stimuli were presented pseudo-randomly.

The visual stimuli were presented on a calibrated (X-Rite calibrator) 24” ViewPixx LCD monitor (1920 x 1080 pixels; 120 Hz refresh rate) placed 54 cm (monkey 1) or 56 cm (monkey 2) from the monkey, using custom software written in MATLAB (Psychophysics Toolbox; Brainard, 1997; Pelli, 1997). Eye position was monitored using an infrared eye tracker (EyeLink 1000; SR Research). Eye position (1000 samples/s), neuronal activity (30,000 samples/s) and the signal from a photodiode was recorded to align neuronal responses to stimulus presentation times (30,000 samples/s) using Blackrock CerePlex hardware.

#### Neural responses

The filtered electrical activity (bandpass 250-5000Hz) was thresholded at 2-3% RMS value for each recording site and the threshold crossing timestamps were saved (along with the raw electrical signal, waveforms at each crossing, and other signals). Spikes were not sorted for these experiments and ‘unit’ refers to the multiunit activity at each recording electrode. The stimulus-evoked firing rate of each V4 unit was calculated based on the spike count responses between 50-250ms after stimulus onset, to account for V4 response latency. The baseline firing rates were calculated based on the spike count responses in the 100ms time-period prior to the onset of the stimulus.

#### Neuron exclusion

For each experimental session, for each unit, the average stimulus-evoked responses across all stimuli were compared with its average baseline activity. The unit was included in further analyses if the average evoked activity was at least 1.1x the baseline activity. This lenient inclusion criterion was chosen because, for the chosen experimental design and stimuli, dimensionality reduced decoding analyses are resilient to noise and benefit from information distributed across many neurons. Each recording experiment yielded data from 90-95 units (mean 94.1).

#### Receptive Field Mapping

A set of 2D closed contours, 3D solid objects, and black-and-white Gabor images were flashed with the same timing as described above in the lower left quadrant of the screen. The positions and sizes were chosen manually across several experiments to home in on the receptive fields of each V4 recording site. Typically, a grid of 5x5 positions and 2 image sizes were chosen such that the images overlapped partially. The spikes were counted within a 50-250ms window after stimulus onset and a RF heat map was constructed for each site. The center of mass of this heat map was chosen as the center of the RF and an ellipse was fit to circumscribe the central two standard deviations. This resulted in centers and extents of the RF of each recording site. The naturalistic image sets for the experiments described below were scaled such that the circular aperture within which the background objects were contained fully overlapped the population RF. This necessitated that the image boundary exceeded the RF of some neurons but the image information outside of the circular aperture was held constant across images.

#### Experiment 1: Effect of task-irrelevant stimulus changes on the ability of V4 neurons to encode a feature of interest about the central object

The first goal of the electrophysiology experiments was to determine if the information about the chosen parameter of the central object (banana position) interferes with information about distracting parameters (background object rotation and depth). To do this, we systematically varied the horizontal position of the object and the background parameters in an uncorrelated fashion. The values and ranges of the object and background parameters were customized for each monkey such that there was a differential response to each condition on average across all other conditions i.e., a 3-way ANOVA for object position and the two background conditions all had a significant main effect (p<0.01). Five values of object position, background depth, and background rotation were chosen and permuted yielding 125 image stimuli. Further details of stimuli can be found in Figure 1 and Figure 1-1, and the associated code and data repositories.

**Figure 1:**
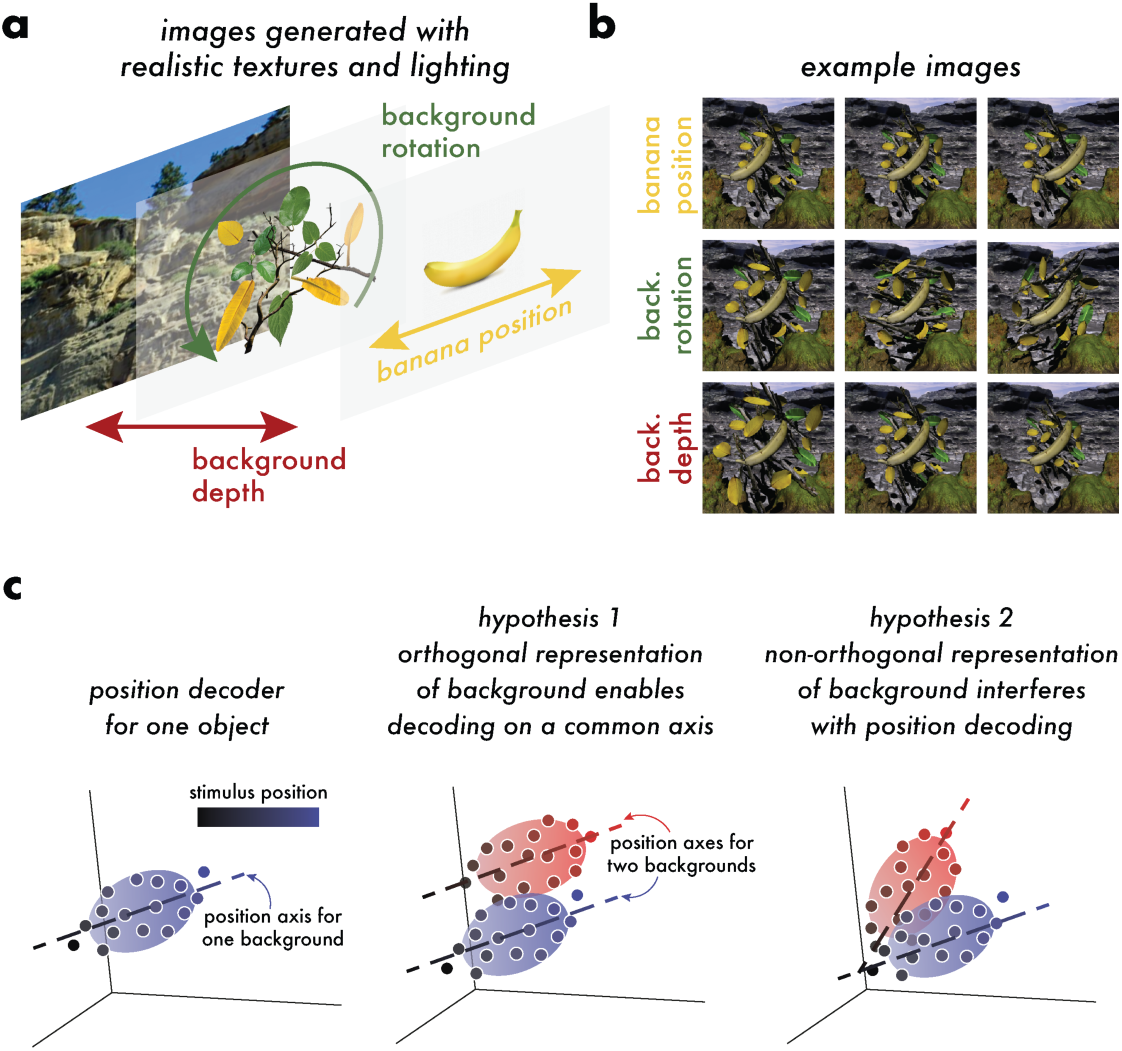
Stimulus design and hypotheses about how neural representations enable generalizable decoding. **a:** We generated photorealistic images with permuted central object (the banana) and background properties using a Blender-based image generation pipeline that gave us control over central- and background-object properties (their position, size, pose, color, depth, luminance, etc.) **b:** Example images showing variations in three parameters – central-object position in the horizontal direction, a rotation of the background objects (leaves and branches), and the depth of the background objects. Five values of each of the three parameters were chosen for each monkey based on receptive field properties (see below), yielding an image set of 5x5x5=125 images. **c:** Hypothesized implications of the neural formatting of visual information on the ability to decode a visual feature. Consider the responses of a population of neurons in a high-dimensional space in which the response of each neuron is one dimension. The population responses to a series of stimuli that differ only in one parameter (e.g. the position of the central object) changes smoothly in this space (left). Responses to a set of stimuli that differ in the same parameter but also have, for example, a difference in the background will trace out a different path in this space (e.g. the red points in the center and right panels). Relative to the first (blue) path, changing the same parameter on a different background could change the population response in a parallel way; more specifically, changing the background could move the population along a dimension that is orthogonal to the dimension encoding the parameter of interest (center). This scenario would enable linear decoding of the parameter of interest that is invariant to changes in the background. Alternatively, the direction that encodes of the parameter of interest could depend on the background (right). Under the linear readout hypothesis, in this case varying the background would impair the ability of a population of neurons to support psychophysical estimation of the parameter of interest.

The data were collected in 26 recording experiments (17 sessions across 11 days from monkey 1, 9 sessions across 8 days from monkey 2). Recording experiments with fewer than 3 repetitions per stimulus image were excluded. Each stimulus was therefore presented between 3 and 16 times yielding between 381-2084 presentations (mean 831).

#### Experiment 2: Relationships between multiple feature dimensions

The second goal of the monkey electrophysiology was to determine different visual features are encoded orthogonally in neuronal population responses. We therefore measured responses to stimuli that varied many features of the central object (banana), including its horizonal position, depth, orientation, and two surface parameters (color and luminance). We also varied the same five features of the background objects (branches and leaves) in an independent way. We used two values of each of the ten features, which we chose to make the ten features equally decodable by the population of V4 neurons (see Figure 5). We therefore measured responses to five repetitions of each of 2^10^=1024 stimuli. Each stimulus image was repeated between two to three times. Because of the large dataset required for this experiment, the data analyzed in Figure 5 were collected from one session from monkey 1.

### Human psychophysics

#### Apparatus

A calibrated LCD color monitor (27-inch NEC MultiSync PA271Q QHD Color Critical Desktop W-LED Monitor with SpectraView Engine; NEC Display Solutions) displayed the stimuli in an otherwise dark room, after participants dark-adapted in the experimental room for a minimum of 5 minutes. The monitor was driven at a pixel resolution of 1920 x 1080, with a refresh rate of 60 Hz and with 8-bit resolution for each RGB channel. The host computer for this monitor was an Apple Macintosh with an Intel Core i7 processor. The head position of each participant was stabilized using a chin cup (Headspot, UHCOTech, Houston, TX). The participant’s eyes were centered horizontally and vertically with respect to the monitor, which was 75 cm from the participant’s eyes. The participant indicated their responses using a Logitech F310 gamepad controller.

#### Stimulus parameters

The entire image subtended 8 degrees in both width and height, the central object subtended ∼4 degrees in the longest dimension, and the circular array of background objects (branches and leaves) subtended ∼5 degrees of visual angle. The images were created using ISET3d at a resolution of 1920 x 1920 with 100 samples per pixel, at 31 equally spaced wavelengths between 400 nm and 700 nm.

#### Psychophysical task

The psychophysical task was a two-interval forced choice task with one stimulus per interval. Each stimulus interval had a duration of 250 ms. Stimuli were presented at the center of the monitor. Between the two stimulus intervals, two masks were shown in succession at the center of the monitor (Figure 5). Each mask was be presented for a duration of 400 ms, for a total interstimulus interval of 800 ms (see Session organization below for mask details). Display times are approximate as the actual display times were quantized by the hardware to integer multiples of the 16.67 ms frame rate.

The task of the participant was to determine whether, compared to the central object presented in the first interval, the central object presented in the second interval was to the left or to the right. Following the two intervals, the participant had an unlimited amount of time to press one of two response buttons on a gamepad to indicate their choice. Feedback was provided via the auditory tones. Trials were be separated by an intertrial interval of approximately 1 second.

The experimental programs can be found in the custom software package Natural Image Thresholds (https://github.com/AmyMNi/NaturalImageThresholds). They were written in MATLAB (MathWorks; Natick, MA) and were based on the software package Virtual World Color Constancy (github.com/BrainardLab/VirtualWorldColorConstancy). They rely on routines from the Psychophysics Toolbox (http://psychtoolbox.org) and mgl (http://justingardner.net/doku.php/mgl/overview).

#### Session organization

The first session experimental session for each participant included participant enrollment procedures (informed consent, vision tests, etc.; see Participants above for details) as well as familiarization trials (see next paragraph) and lasted one and a half hours. The additional experimental sessions lasted approximately one hour each.

For the first session only, the participant began with 30 familiarization trials. The familiarization trials comprised, in order: 10 randomly selected easy trials (the largest position-change comparisons), 10 randomly selected medium-difficulty trials (the 4th and 5th largest position-change comparisons), and 10 randomly selected trials from all possible position-change comparisons. The familiarization trials did not include any task-irrelevant variability and data from these trials was not saved.

In each session, there were two reference positions for the banana, and for each reference position there were 11 comparison positions: five comparison positions in the positive horizontal direction, five comparison positions in the negative horizontal direction, and a comparison position of 0 indicating no change. On each trial, one interval contained one of the two reference stimuli and the other interval will contain one of that reference stimulus’s comparison stimuli. The order in which these two stimuli were presented within a trial was selected randomly per trial.

A block of trials consisted of presentation of the 11 comparison positions for each of the two reference positions for a total of 22 trials per block. The trials within a block were run in randomized order. Each was completed before the next block began. Each block was repeated 7 times in a run of trials, for a total of 154 trials per run.

Within each run of 154 trials, a single background variation condition was studied. There were three such conditions, as described in more detail below – “no variation”, “rotation only”, and “rotation and depth”. Two runs for each of the three conditions was completed in each experimental session, and except as noted in the results, each subject completed 6 sessions. The six runs were conducted in random order within each session, and each run was separated by a break that lasted at least one minute and during which the participant was encouraged to stand or stretch as needed. After a minimum of one minute, the next run was initiated when the participant was ready.

Additionally, each session began with four practice trials (including in the first experimental session, where these practice trials were preceded by the familiarization trials as described). Each run after the first also started with one practice trial. The practice trials were all easy trials as described above and not include any task-irrelevant variability. The data from the practice trials was be saved. The maximum variation in background features was matched to the maximum variation in the neurophysiology experiments but sampled more finely as described for each of the variation blocks below.

For the “no variation” condition, there were not any changes to the background objects (the branches and leaves). This run determines the participant’s threshold for discriminating the horizontal position of the central object without any task-irrelevant stimulus variation.

The “rotation only” run introduced task-irrelevant variability single task-irrelevant feature: rotation of the background objects. For each trial, a single rotation amount was drawn randomly from a pool of 51 rotations, and the background objects (leaves and sticks) in the stimulus were all rotated by that amount around their own centers. The rotation was drawn separately (randomly with replacement) for each of the two stimuli presented on a trial (the reference position stimulus and the comparison position stimulus). Thus subjects had to judge the position of the central object across a change in the background, so that any effect of background variation on the positional representation of the central object would be expected to elevate threshold. The pool of 51 rotations comprised: a rotation of zero (no change to the background objects), 25 equally spaced rotations in the clockwise direction in 2-degree intervals, and 25 equally spaced rotation amounts in the counterclockwise direction in 2-degree intervals.

“Rotation and depth” runs had variation in two task-irrelevant features: rotation and depth of the background objects. For this run, there was a pool of 51 rotations, but along with the rotation of the background objects, these objects also varied in depth. There were 51 possible depth amounts (one depth amount of zero, 25 equally spaced depth amounts in the positive depth direction, and 25 equally spaced depth amounts in the negative direction; depth amounts ranged from -500 mm to 500 mm in the rendering scene space). One of the of images was a rotation of zero and a depth amount of zero. For the remaining 50 images in the pool, each of the remaining 50 rotation amounts was randomly assigned (without replacement) to one of the remaining 50 depth amounts. The same depth shift was applied to each of the background objects. From this pool of 51 images, a single image was randomly drawn (with replacement) for each of the two stimuli presented in the trial.

Finally, as noted above (see Psychophysical task), two masks were shown per trial during the interstimulus interval. All masks across all background variation conditions were created from the same distribution of stimuli (stimuli with “no variation”, thus containing no task-irrelevant noise). To create each of the two masks, first the central object positions in the first and second intervals of the trial were be determined. The two stimuli with that matched the central object positions in the first and second intervals were then used to create the trial masks. For each of these two stimuli, the average intensity was calculated in each RGB channel per 16 x 16 block of the stimulus. Next, each 16 x 16 block of a mask was randomly drawn from the mask corresponding to the two stimuli. Thus, the two masks shown per trial were each a random mixture of 16 x 16 blocks from stimuli with the two central object positions for that trial.

### Statistical Analysis and Quantification

#### Monkey electrophysiology

##### Cross-validated Parameter Decoding (Figure 2)

First, the response matrix (multiunit spike rates for each site for each image stimulus presentation) was reduced to 10 dimensions of activity. This ensured sufficient dimensionality for the decoding of object and background parameters and explained between 87.8% and 94.8% (mean 91.2%) of the variance across stimulus responses. (Parameter decoding without dimensionality reduction produced qualitatively similar results.) Then, for each background condition (unique combination of background rotation and depth – or “specific decoding”), the object position in each presentation/trial was decoded from neural responses by learning regression weights from all other trials (leave-one-out cross-validation). The same procedure was repeated for “general decoding” where the background parameters were ignored (Figure 2b).

**Figure 2:**
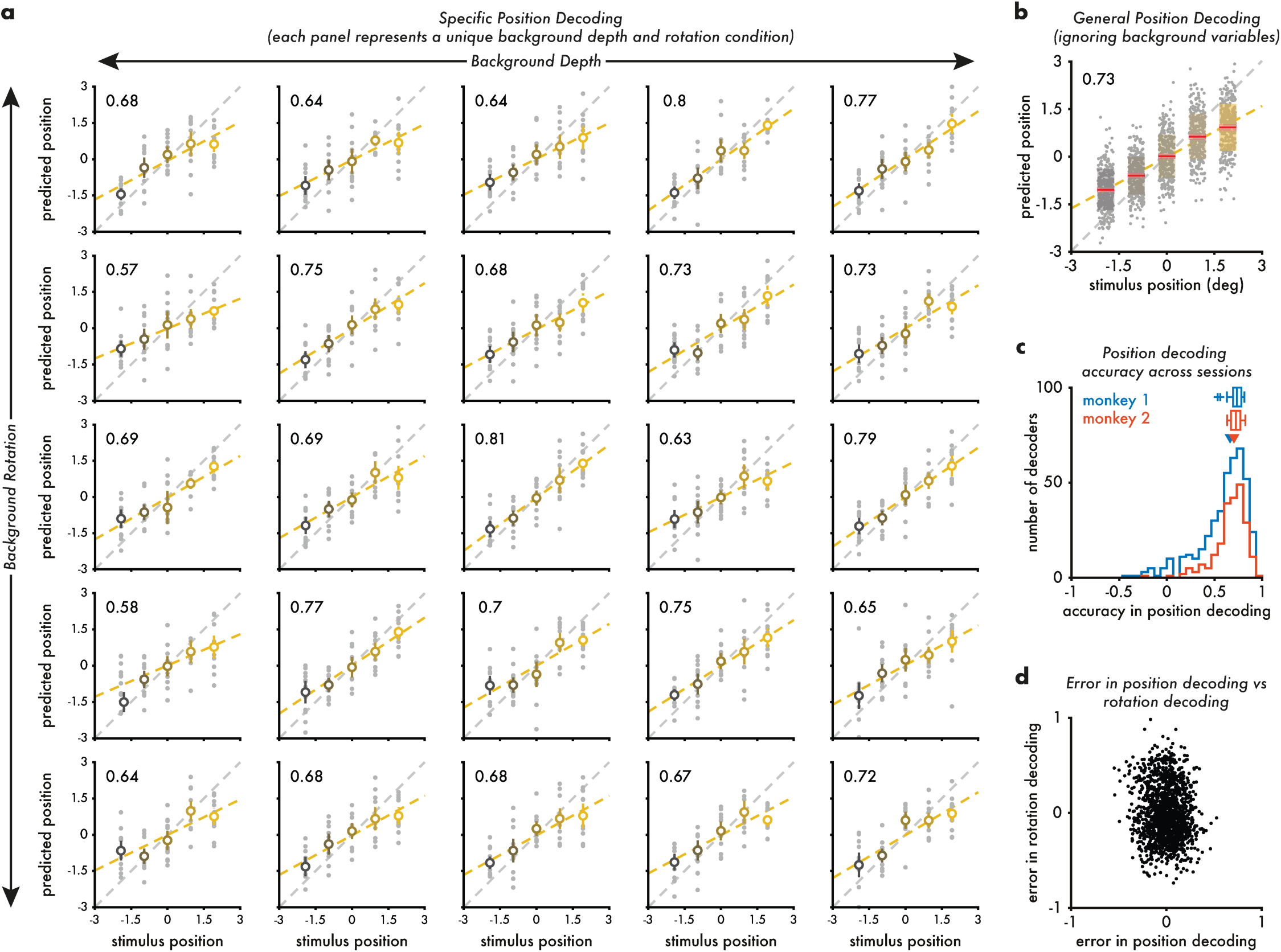
Object position decoding from V4 population responses is consistent across background variations. **a:** We can linearly decode object position for each background stimulus (for the example session shown here). Each panel represents a unique configuration of background rotation and depth, with rows representing variations in rotation and columns representing variations in depth. Each gray point shows the decoded position for a single image presentation in this session. These points depict the actual object position (x-axis, in visual degrees relative to the center of the image) and the decoded position (y-axis) using a separate, cross-validated linear decoder for each unique background. The open circles represent the trial-averaged predicted position (vertical length is the standard deviation). The number in the top-left is the correlation between the actual and decoded positions and the yellow dashed line is a linear fit. Gray to yellow gradient is a redundant cue for stimulus position variation. The gray dashed line represents the identity. **b:** Position decoding is largely consistent across background variations. This plot is in the same format as those in A. Here, a common decoder that ignores variations in the background and therefore incorporates all stimulus presentations is used. The data are the same as those shown in A. Compare with Figure 2-1a and 2-1c for background rotation and depth decoding. **d:** Distribution of specific decoder accuracies (correlation) across all sessions for each monkey (each session contributed 25 values to the histogram). Blue and red arrows represent the median accuracy (0.662 for monkey 1, 0.703 for monkey 2). The box plots above the histograms summarize general decoder accuracy across sessions for each monkey. The central line indicates the median (0.735 for monkey 1, 0.724 for monkey 2), box edges indicate 25 and 75 percentiles, whiskers indicate minimum and maximum values, and + symbols indicate outliers. Compare with Figure 2-1b and 2-1d for background rotation and depth decoding. **d:** Error in decoding object position (across background variations) for each trial compared with the error in decoding background rotation. See Figure 2-1e for comparison with error in trial-wise background depth decoding.

Decoding accuracy was defined as the correlation between the actual values and the decoded values. Perfect decoding would result in an accuracy of 1 and chance decoding in an accuracy of 0. We did not encounter decoding accuracies below 0. We also calculated other decoding performance measures like mean squared error, cosine distance, etc. While other measures provide more sensitivity in the specific kinds of decoding error, their estimate of aggregate performance was qualitatively similar to correlation-based measures. Error in decoding was defined as the difference between the predicted object position and the actual position (Figure 2d). The same procedure for specific and general decoding was repeated for each of the two background conditions as well (Figure 2-1).

#### Angle calculation (Figure 3)

To calculate the angle between the specific decoders, an n-dimensional line was fit to the dimensionality reduced responses and the unit vector was found. The angle between each specific decoder and the decoder for the central condition was calculated as the arc-cos of the dot product of the two unit vectors.

**Figure 3:**
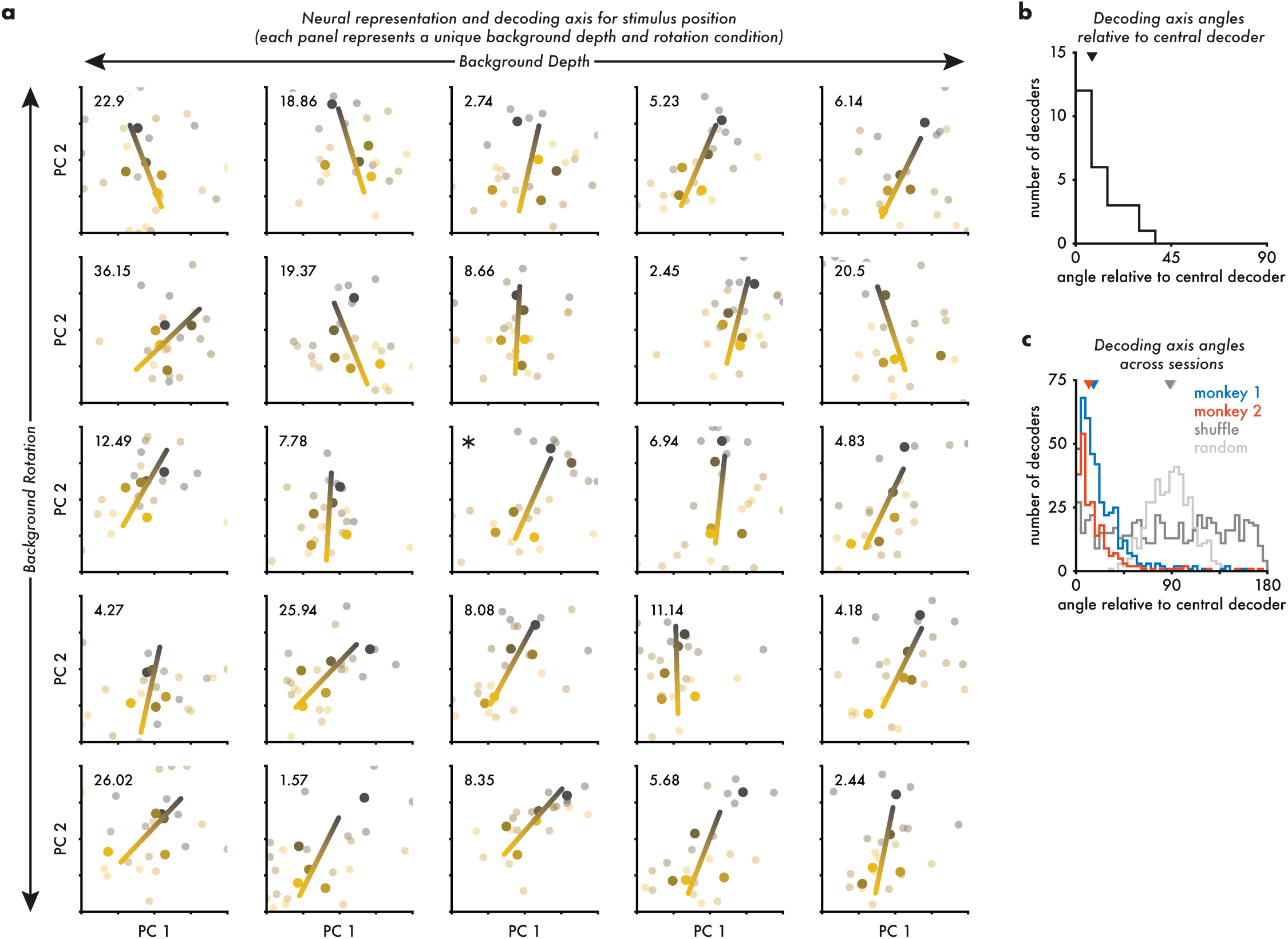
Object position axes across background variations are aligned with each other. Since object position decoding is tolerant to background variations, we tested whether the linear decoding axes for each background configuration were aligned by visualizing the decoders in the first two principal components of the neural response space. These dimensions were computed for the full set of neural responses obtained in each session. **a:** As with Figure 2a, each panel represents a unique configuration of the background rotation and depth with rows representing variations in rotation and columns representing variations in depth. Each dim point represents a single image presentation, and bright points represent trial-averaged responses. Gray-to-yellow gradient represents monotonic variation in object position. A gradient line was fit to the responses for each background condition, shown here in two dimensions for illustration. The lines shown have been normalized to have the same length in the projected space shown. The text label at the top left represents the relative angle between each decoder and the central decoder (the middle background condition plot, marked with ∗) calculated in the full dimensional space of responses used for decoding. **b:** Distribution of angles in A as a histogram. Arrow at the top represents the median angle for this session (7.78°). **c:** Distribution of relative decoder angles across all object position decoders (like those in A) across sessions for both monkeys (blue and red distributions). Blue and red arrows represent the median of angles across sessions (16.4° for monkey 1, 12.07° for monkey 2). Dark gray distribution represents the angles of object position decoders after shuffling the position values for each trial (median 88.13°, shown as dark gray arrow). Light gray distribution represents the angles between randomly chosen vectors of the same dimensionality as the neural population space (median 89.99°, shown as light gray arrow).

#### Linear discriminant analysis and comparison with human psychophysics (Figure 4c)

To directly compare human psychophysics discrimination accuracy with decoding results, we matched the three blocked conditions – no background variation, rotation only, and rotation and depth variation – by subsampling trials from the 5x5x5 stimulus set from experiment 1. For the three conditions, we either found all pairs of trials, pairs of trials that varied in rotation only (by holding background depth at the central value), or pairs of trials that varied in depth only (by holding background rotation at the central value). Then, for 200 folds, we sampled a maximum of 500 pairs of trials and depending upon the object position on those trials, we assigned a left or right choice. If the positions were identical, we randomly assigned the choice for that pair. We then collated the responses across the pairs of trials and a fit linear discriminant in a leave-one-out fashion to predict the correct choice. The classification prediction accuracy for each of the three blocked conditions was calculated independently.

**Figure 4:**
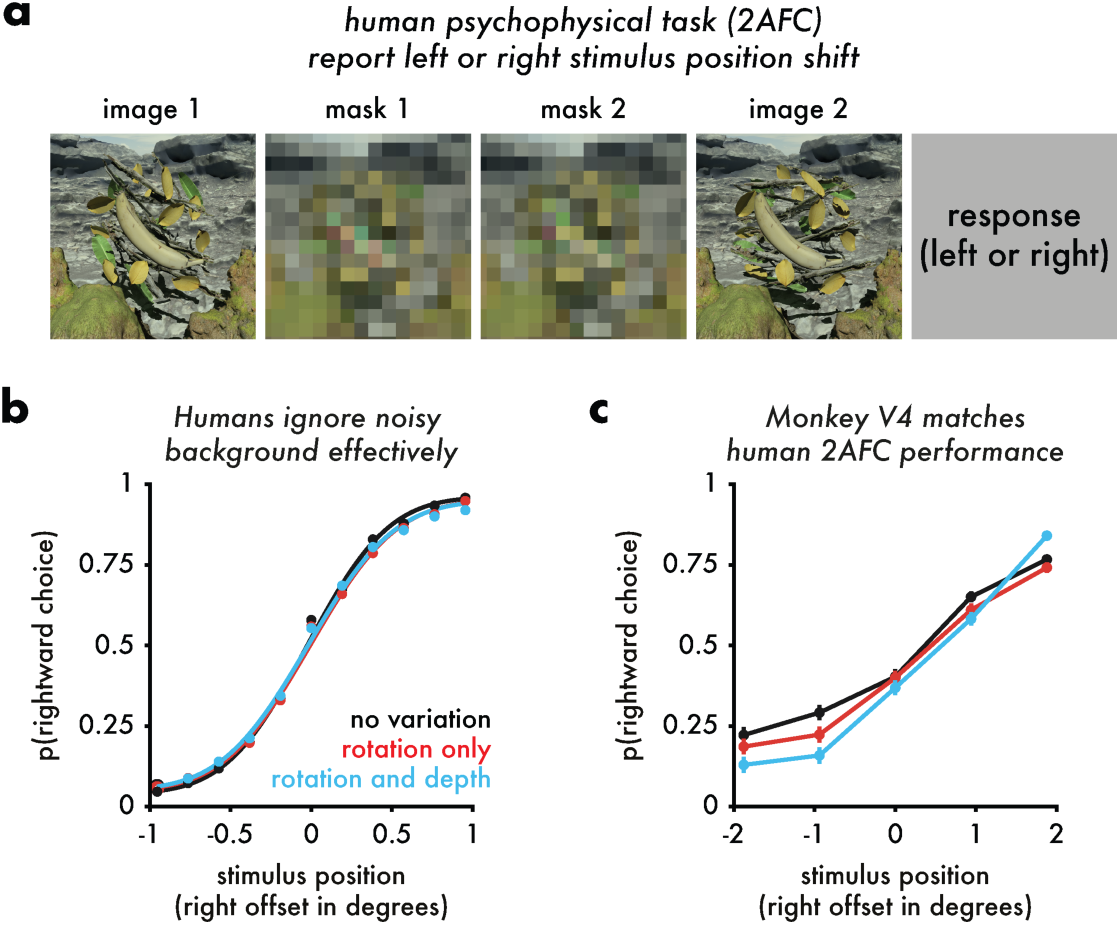
Human psychophysics experiments suggest that discrimination of object position is unaffected by variation in the stimulus background. Since object position representation in V4 neurons is robust to background variation, we tested whether changing background properties affects the decoding of object position in humans. **a:** Human psychophysical task. Two images containing a banana were presented, with two masks in between. Participants were instructed to report whether the relative position of the banana in image 2 was left or right of that in image 1. In blocks, background rotation and/or depth were held constant or varied as described below. **b:** Averaged (across participants, N = 10) psychometric functions for each background variation condition (individual participant performance shown in Figure 4-1). Three background conditions were tested: black: no background variation between the two presentations of the banana on each trial; red: background rotation changed randomly across the two presentations of the banana, but background depth was held fixed; blue: both background rotation and depth were randomized across the two presentations of the banana. Across participants, object position change detection performance was not substantially different across the three background variation conditions. **c:** To compare human behavior and monkey electrophysiology, we selected stimulus presentations in the monkey experiments to approximate the three background variation levels used in the human psychophysics experiment (see Methods for details of sample matching). We trained linear discriminants to separate trials into right or left position shift, for each background variation condition. During training, the trials with the banana in the central position were randomly assigned to be left or right. Classifier performance also did not differ substantially across the background variation conditions.

#### Cross-decoding analysis (Figure 5)

For experiment 2, even though only two values were chosen for each of the five object and five background parameters, linear regression was chosen to instead of classification using discriminant analysis for comparison with decoding analyses in the previous experiment. Even though each stimulus image was only repeated 2-3 times, since each parameter could take one of two values each, all unique pairs of images would be informative about at least one parameter change. To enable cross-decoding, we altered the cross-validation procedure. For each parameter pair, for each of 100 folds, we randomly split all image presentations evenly into training and testing sets (uneven splits also produced qualitatively similar results). We then trained a linear regression model for one parameter using the training trials and used it to predict the values of the other parameter for the held-out testing trials. The decoding accuracy was calculated as the average correlation across folds between the actual and decoded parameter values. Since each parameter decoder was trained while ignoring all other parameter variations, the diagonals in Figure 5b are akin to the general decoder accuracy for those parameters, and the off diagonals correspond to how well those general decoders are aligned to the representations of the other parameters. The diagonal correlations were all significantly above 0 (p < 10^-80^; t-test across folds) and none of the off-diagonal correlations were except the cross-decoding of background and object color.

### Human psychophysics (Figure 4a-b)

Per session, the participant’s threshold for discriminating object position was measured for each background variation condition. First, for each comparison position, the proportion of trials on which the participant responded that the comparison stimulus was located to the right of the reference stimulus was calculated. Next, the proportion the comparison was chosen as rightwards was fit with a cumulative normal function using the Palamedes Toolbox (http://www.palamedestoolbox.org). To estimate all four parameters of the psychometric function (threshold, slope, lapse rate, and guess rate), the lapse rate was constrained to be equal to the guess rate and to be in the range [0, 0.05] and the maximum likelihood fit determined. Threshold was calculated as the difference between the stimulus levels at performances (proportion the comparison was chosen as rightwards) equal to 0.7602 and 0.5 as determined by the cumulative normal fit.

## Results

### Central hypothesis: orthogonal representations enable observers to ignore irrelevant visual information

We tested the hypothesis that task-irrelevant information will not affect a perceptual judgment if the representations of the task-relevant and irrelevant features are orthogonal (Hong et al., 2016). Figure 1a-b depict how we used computer graphics to parametrically vary different scene features, such as the position of a central banana, the rotational position of objects in the background, and the depth position of objects in the background. Using this set of stimulus variations, consider the effect of varying the position of the banana on a hypothetical neural population response as illustrated in the left panel of Figure 1c. Each point in the plot represents the noisy population response to one presentation of an image, illustrating how varying banana position against a fixed background can trace out a line in the high-dimensional neural population space. For this background, the position of the background could be read out by projecting the population response onto the line shown (labeled ’position axis for one background’ in the figure). The middle panel of 1c shows a way that varying the background objects could affect this line in an orthogonal manner. Here the line tracing out the neural population response to the banana at various positions is shifted in a direction orthogonal to the position axis shown in the left panel.

Although a different line is swept out by varying banana position against this second background, projecting onto the line for the first background continues to accurately decode the banana position. If, on the other hand, changing the background causes a change in the position axis that is not orthogonal (right panel of Figure 1c), projecting onto the line for the first background will not provide an accurate linear position readout. Thus we test the hypothesis that changing an irrelevant feature of the background (e.g. the position of background objects) will not impact perception of the task-relevant feature if the irrelevant background changes are orthogonal to the relevant ones (Figure 1c middle).

### Naturalistic stimuli with parameterizable properties

To test these predictions, we created naturalistic stimuli that had many parameterizable features (Figure 1a-b). We parametrically varied the position of a central object (banana), and the rotation and depth of background objects (leaves and branches) set against a larger fixed contextual scene (rocks, moss-covered stumps, mountains, and skyline). We presented these stimuli within the joint receptive fields of recorded V4 neurons (Figure 1-1) while each of two monkeys fixated on a central point. In sum, we recorded V4 responses in 26 experimental sessions across two animals (85-94 visually responsive multiunits per session). Most of the units in our measured population were modulated by the position of the central object and the variations in background rotation and depth (Figure 1-1d).

### V4 neurons robustly encode stimulus position for each stimulus background

We first measured the extent to which V4 neurons encode banana position by linearly decoding that position for each unique background stimulus. Figure 2 shows that for each unique background configuration (rotation and depth), V4 neurons from a single session support good linear decoding of the position of the banana (each of the 25 panels in Figure 2a is for a specific background configuration; each gray point in the panels shows decoded banana position for a single presentation; the mean and standard deviation for each position - open circles and error bars - summarize our ability to predict the position of the banana from the activity of V4 neurons). The numbers at the upper left of each panel provide the correlation between the predicted and actual banana stimulus position (mean performance = 0.698).

### V4 representations of stimulus position and background features are approximately orthogonal

Three lines of evidence support the view that the neural representation of the position of the central object (banana) in V4 is orthogonal to the representations of features of the background (depth and rotation of the leaves and branches) in our stimuli.

First, we compared our ability to decode stimulus position in each background (Figure 2a) with our ability to decode the position of the banana across the entire stimulus set (Figure 2b), where the variability in terms of background depth and rotation has the opportunity to intrude. The ability of a single decoder to read out banana position across all background variations is similar to that of the decoders that were optimized for each unique background stimulus (compare Figure 2b with 2a) and is high across all sessions for two monkeys (Figure 2c). We also found that our ability to decode background rotation (Figure 2-1a and 2-1c) and depth (Figure 2-1b and 2-1d) was similar using a single decoder for all stimuli and using a unique decoder for each combination of the other two features, suggesting that the same population of V4 neurons encodes all three axes of image variation well and orthogonally.

Second, we found that on a trial-by-trial basis, errors in the decoded estimates of banana position are not correlated with errors in decoding of background rotation and depth (Figure 2d and Figure 2-1e). This lack of correlation also suggests that the representations of banana position are independent in V4 from representations of background rotation and depth.

Finally, if the representations of stimulus position and background parameters are orthogonal, then the decoders optimized for each unique stimulus (Figure 2a) should be mutually aligned in neural population space. Put another way, if the representation of stimulus position is robust to variation in the background, this representation should vary along the same direction across backgrounds. To probe this, we calculated the line in neural population space that best explains population responses to each stimulus position for each background. These are depicted in Figure 3a for each unique background condition (plotted for the first two principal components of the population neural responses for visualization purposes only; each point is a trial, each bright point is the average population response to a particular object position, and gray to yellow point colors represent the five object position values). The angle in the population space between the decoders for each unique background and the decoder for the background configuration whose decoder is shown in the center of 3a marked with ∗ (chosen as a reference simply to define the origin of the angular measure) is indicated in degrees. The distribution of angles for this example session (Figure 3b) and across all sessions in both animals (Figure 3c) is skewed toward much smaller angles than expected by chance – gray distributions in Figure 3c depicting a median of ∼90° for decoders trained on shuffled (randomizing trial labels within background configuration) responses to each unique background condition (labeled “shuffle”; dark gray) and angles between random vectors in a response space with the same dimensionality as the neural decoding space (labeled “random”; light gray). Together, these recording results support the idea that the neural representation of stimulus position is orthogonal to the representations of background rotation and depth in V4.

### Human subjects discriminate stimulus position robustly with respect to background variation

A prediction of our central hypothesis is that when the representations of two stimulus features are orthogonal in the brain, varying one should not impact the ability of subjects to discriminate the other. We tested this hypothesis by measuring the ability of human observers to discriminate the position of the central banana in our stimuli amid variation in the background rotation and depth. By using a threshold paradigm, we test this idea for stimulus step sizes that approach the limits of perception.

Human subjects viewed two images of the banana separated by two different masks (Figure 4a) and reported whether the banana in the second image was positioned to the left or right of the banana in the first presentation. The offset between the two banana positions was varied systematically, to allow determination of discrimination threshold. Across blocks of trials, we varied the amount of within-trial image-to-image variability in the background objects across the two presentations of the banana. When there was background variation, intrusion of that variation on decoding of banana position would manifest itself as an elevated discrimination threshold if the representation of the banana position was not orthogonal to that of the background features(Singh et al., 2022; Reynolds and Singh, 2023). However, consistent with the idea that the orthogonal representations of object position and background features that we found in the neural recordings enables background-independent perception, introducing variability into background did not significantly impact the position discrimination performance of human subjects (Figure 4b and Figure 4-1). To make a direct comparison between parameter decoding of neural representations and human psychophysical performance, we partitioned neural data into “no variation”, “rotation variation only”, and “depth and rotation variation” groups and trained linear discriminants to classify left or right position difference between pairs of presentations (Figure 4c). The cross-validated discrimination performance of these classifiers also did not differ across the three background variation groups, as with the human psychophysical performance. Thresholds for the neural classifiers are higher than those for the human subjects (note difference in x-axis scale between Figure 4b and 4c), but this is not surprising given the difference in visual field location and the fact that it seems unlikely that we recorded from all of the neurons that support psychophysical performance.

### At least ten object and background features are represented approximately orthogonally in V4

To test the extent to which the orthogonality of representations of different features generalizes to other features of the banana and background in our stimuli, we measured V4 responses to a large image set where the color, luminance, position, rotation, and depth of both the background and object each took one of two values (this yields 2^10^ = 1,024 unique images; Figure 5a). If any two parameters are encoded orthogonally in neural population space, then it should be possible to linearly decode those parameters successfully despite the variation in the others. Conversely, a decoder trained on one parameter should not provide information about the others. To test these predictions, we trained linear decoders for each of the object or background features and then tested our ability to decode each of the ten features with each decoder.

Each of the ten features was encoded in the V4 population despite the variation in the other features, meaning that the correlation between the actual value of the feature parameter and the value predicted by a cross validated linear decoder was above chance (diagonals in Figure 5b). In addition, the correlation between a given parameter and the value predicted by a decoder trained on a different parameter (off-diagonals in Figure 5b) was indistinguishable from chance except in one case (the color of the central object and background objects). These observations suggest that a population of V4 neurons can encode a relatively large number of natural scene parameters independently, enabling observers to avoid distraction by task-irrelevant stimulus features. The observation that there is an interaction between central and background objects presents an opportunity for future work to test the prediction that task-irrelevant variation in background object color should affect psychophysical discrimination of central object color, an outcome that would be consistent with the results of Singh et al. 2022.

**Figure 5:**
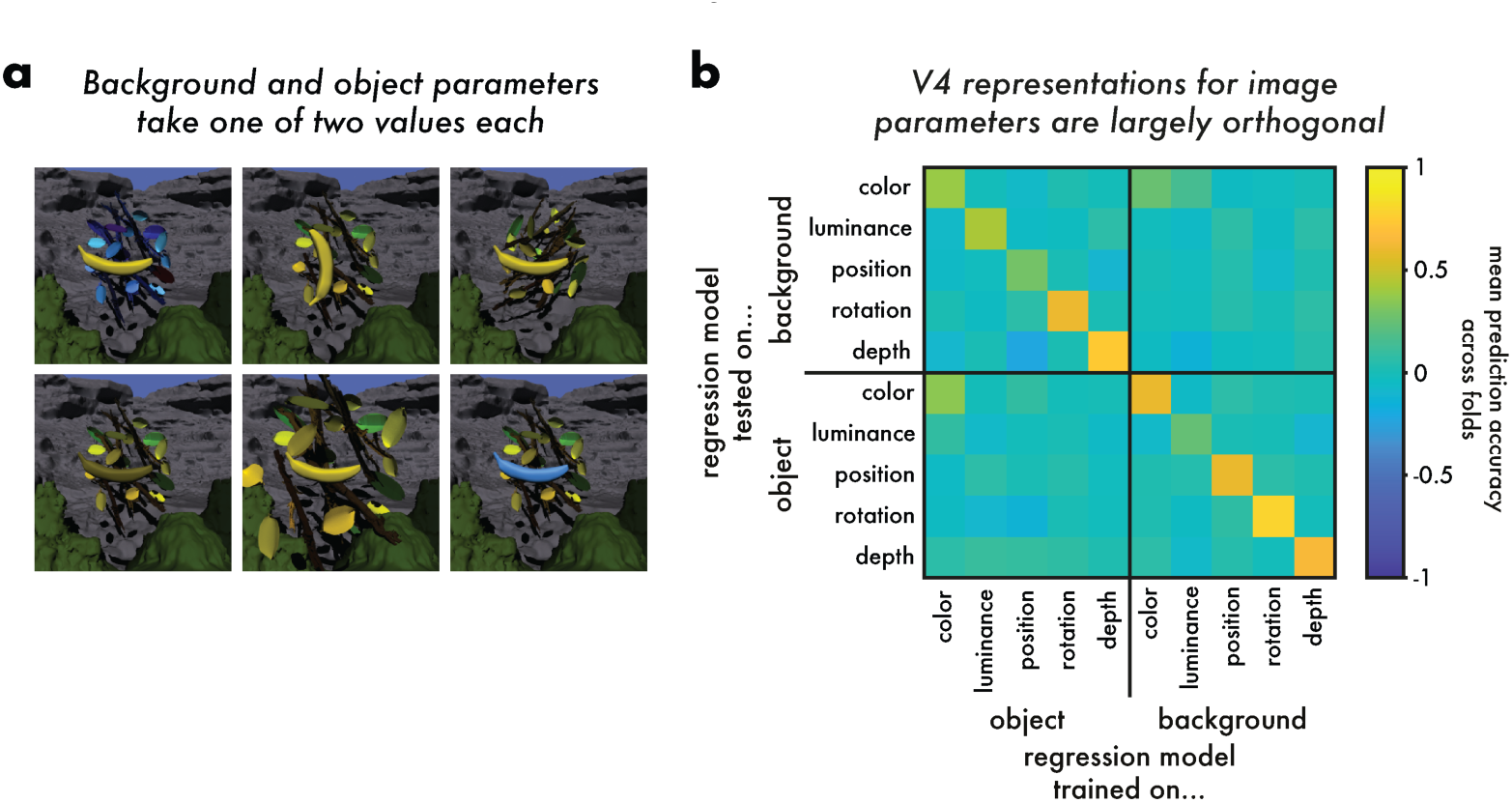
Orthogonal representations for variations in up to 10 object and background features. We generated a large image set where the color, luminance, position, rotation, and depth of both the background and object each took one of two values yielding 2^10^ = 1,024 images. We collected V4 population responses to these images as in Figure 2c. **a:** Example images illustrating object and background parameter variation. **b:** If two features are encoded orthogonally (independently) in neural population space, then a decoder trained on one feature should not support decoding of the other feature. We trained linear decoders of V4 responses for each of the object or background features (x-axis) and tested the ability to decode each of the 10 features. The diagonal entries provide, for each feature the correlation between the decoded and actual feature parameter values for a decoder trained on that feature. Correlations obtained through cross-validation. Decoding performance was above chance (correlation of 0) for all features (p < 10^-80^; t-test across folds). The off-diagonal values depict evaluate performance of a classifier trained on one feature (x-axis) for decoding another (y-axis). This cross-decoding performance was not distinguishable from chance except in the case of color of the object and background

## Discussion

Using a combination of multi-neuron electrophysiology in monkeys and human psychophysics, we tested the hypothesis that features of irrelevant objects in naturalistic scenes will not interfere with the perception of target object features when their representations are orthogonal in visual cortex. We demonstrated that 1) in monkey area V4, the representation of object position is orthogonal to the representations of many irrelevant features of that object and the background, and 2) consistent with our hypothesis, threshold for human observers to judge a change in object position was unaffected by the variations in the background stimulus that were shown neuronally to have orthogonal representations in monkey V4.

### Relationship to the notion of untangling and representational geometry

The conditions under which objects can be disambiguated from neural population responses have been studied using the concept of *untangling* (DiCarlo and Cox, 2007; Rust and Dicarlo, 2010; DiCarlo et al., 2012; Pagan et al., 2013). Untangling has been primarily discussed in the context of object classification. The hypothesis is that different objects can be appropriately classified (e.g. discriminating images of bananas from images of leaves) when the neural population representations of those objects are linearly separable in the face of irrelevant variations in the images (e.g. changes in position, orientation, size, or background). Support for this hypothesis comes from the observation that as one moves from early to late stages of the primate ventral visual stream, representations of different object categories become more linearly separable (DiCarlo et al., 2012; Yamins et al., 2014; Majaj et al., 2015; Hong et al., 2016; Hénaff et al., 2019). Progress has been made in understanding how the tuning functions and mixed selectivities of neurons support untangled population representations (Rigotti et al., 2013; Fusi et al., 2016; Kriegeskorte and Wei, 2021).

The untangling framework has been extended to address the structure of neural populations that represent object categories more generally by characterizing the geometry of the high dimensional representational manifolds (Yuste, 2015; Chung et al., 2018; Saxena and Cunningham, 2019; Cohen et al., 2020; Chung and Abbott, 2021; Jazayeri and Ostojic, 2021). The capacity and dynamics of representational geometries in visual cortex correlate with classification behavior (DiCarlo et al., 2012; Rajalingham et al., 2018; Stringer et al., 2019), in parietal and prefrontal cortices with perceptual decision making (Mante et al., 2013; Bernardi et al., 2020; Okazawa et al., 2021b; Ehrlich and Murray, 2022), and in motor cortex with control of muscle activity (Shenoy et al., 2013; Sussillo et al., 2015; Gallego et al., 2018; Russo et al., 2018).

The untangling framework has also been extended to the linear readout of object properties from orthogonal neuronal representations (Hong et al., 2016). Here, we employ this extension to explore the ability of observers to represent the continuous values taken on by lower-level visual features like the ones we studied. To connect the untangling idea as applied to object classification with the formulation here, note that the linear separability employed for object-class discrimination is effective when the effect of irrelevant image variations is orthogonal to the hyperplane that separates the classes in the neural population space. Thus we use the same concept of orthogonality but apply it to linear decoding of feature parameter values rather than to linear separation of object classes. Additionally, we vary the irrelevant features (background depth and rotation) parametrically to estimate independent position axes and demonstrate that across irrelevant feature variation, the position encoding axes are parallel to each other. We link the neural orthogonality to behavioral performance by deploying a recently developed psychophysical paradigm (Singh et al., 2022; Kramer et al., 2023; Reynolds and Singh, 2023) to quantify the degree of perceptual orthogonality between the same task-relevant and task-irrelevant stimulus features that we study neurally.

Other studies consistent with this line of thinking have also considered features and have demonstrated that neural responses to relevant and distracting features of simple stimuli are linearly separable in the brain areas (or analogous layers of deep network models of vision) that are thought to mediate that aspect of vision (DiCarlo and Cox, 2007; Khaligh-Razavi and Kriegeskorte, 2014; Yamins et al., 2014; Chung et al., 2016; Cohen et al., 2020). Indeed, a previous study in our lab also found a relationship between our ability to linearly decode visual information the activity of neural populations in monkeys and the ability of human observers to discriminate the same stimuli(Kramer et al., 2023). Our conclusions are also consistent with those reached in a study that considered neuronal representations in V4 and IT and behavioral estimates of the properties of objects presented against natural image backgrounds (Hong et al., 2016). That study found increasing orthogonality of representation from V4 to IT, and good behavioral estimation of the orthogonally represented properties. Given those results, it seems possible that our stimuli would have revealed increased orthogonality in areas further along the processing hierarchy than the V4 site of our electrode array; such a result would not change the general conclusions we draw about the relation between orthogonality and behavior performance.

### Opportunities from studying parameterizable naturalistic images

The present study extends the measurements of the relationship between the neural untangling of lower-level features and visually guided behavior with respect to features in naturalistic images. Our view is that the perception of object features in complex natural images provides increased power for testing the untangling hypothesis in the context of feature decoding. Unlike the case of simpler stimuli, the number of task-irrelevant features available for manipulation is larger and is likely to more fully challenge the coding capacity of neural populations whose representations are of limited dimensionality (Chung and Abbott, 2021). Furthermore, visual distractors (like variation in the background) heavily influence scene categorization performance in artificial stimuli but not natural stimuli, suggesting that orthogonal feature representations in natural stimuli are more resilient to noise (Zhou et al., 2000; Chung et al., 2018). Studying the relationship between neurons and visually guided behavior using parameterizable naturalistic images solves many of the challenges inherent in using simple artificial stimuli on the one hand or natural images on the other (Felsen and Dan, 2005; Rust and Movshon, 2005; Martinez-Garcia et al., 2019; Cowley et al., 2023; Ding et al., 2023; Maheswaranathan et al., 2023). The graphics-generated stimuli we employ strike a balance between the experimental control available through parameterization and the ability to measure principles governing neural responses to and perception of features of natural images that are difficult or impossible to glean using artificial stimuli.

### Opportunities from cross-species investigations of visual perception

Our results highlight the power of pairing neural population recordings in animals with behavior in humans for understanding the neural basis of visual perception. Although simultaneously recording neurons and measuring behavior has many advantages, comparison with human performance provides some assurance that the neural results obtained in an animal model generalize to humans. In addition, our approach links observations from the more peripheral visual field locations where for technical reasons the neural recordings are most often made, to the central visual field locations that are typically the focus of studies with human subjects.

Since the monkeys were simply rewarded for fixating during the recordings, our experiments focus on neural population activity that is stimulus driven, rather than reflecting internally driven processes like attention or motivation. In future work, it will be interesting to merge our knowledge of how stimulus-driven and internal processes combine to influence neuronal responses and performance on visual tasks.

### Mechanisms supporting orthogonality

Our study quantifies how neural populations represent multiple naturalistic stimulus variations, but it does not provide direct insight about how the encoding and processing of visual stimuli produce those representations. Under biologically realistic assumptions, simulations show that although it is possible to learn about the orthogonality of feature representations within a population from small population recordings, it is generally not possible to characterize the role of each recorded neuron (Ruff et al., 2018b).

In recent years, a large number of studies have demonstrated that neural networks trained to categorize natural images produce representations that bear strong resemblance to neural representations in the ventral visual stream (Oleskiw et al., 2018; Pospisil et al., 2018; Bashivan et al., 2019; Srinath et al., 2021; Cowley et al., 2023). These models provide an opportunity to understand the conditions under which aspects of natural stimuli are represented orthogonally, which is a subject of ongoing work (Majaj et al., 2015; Chung et al., 2016, 2018; Hong et al., 2016; Cohen et al., 2020; Ni et al., 2022; Kramer et al., 2023). We hope that our results will lead to a productive coupling of computational analysis of the mechanisms by which orthogonal representations emerge with behavioral experiments using the same parametrically varied computer-graphics stimuli.

## Conclusion

Our results provide behavioral and neurophysiological evidence supporting the powerful untangling hypothesis, further extend the study of untangling to representations of features of objects and backgrounds and demonstrate the value of parameterizable naturalistic images for studying the neural basis of visual perception. They also suggest a promising future of investigating the neural basis of perceptual and cognitive phenomena by leveraging the complementary strengths of multiple species.

## Acknowledgements

We are grateful to K. McKracken for providing technical assistance, to Douglas Ruff, Cheng Xue, and Lily Kramer for comments on an earlier version of this manuscript and helpful comments and suggestions regarding data analysis. This work is supported by Eric and Wendy Schmidt AI in Science Postdoctoral Fellowship (to R.S.), the Simons Foundation (Simons Collaboration on the Global Brain award 542961SPI to M.R.C, postdoctoral fellowship to A.M.N.), the National Institutes of Health (awards R01EY022930, R01EY034723, and RF1NS121913 to M.R.C, 1K99NS118117-01 to A.M.N, K99EY035362 to R.S.)

## Author Contributions

Conceptualization - A.M.N., M.R.C., D.H.B.; Methodology - R.S., A.M.N., M.R.C., D.H.B.; Software - R.S., A.M.N., D.H.B; Formal Analysis - R.S., A.M.N., C.M. M.R.C., D.H.B.; Investigation - R.S., A.M.N., C.M.; Data curation - R.S., A.M.N., C.M.; Writing – Original Draft - R.S., M.R.C., D.H.B. ; Writing – Review & Editing - R.S., M.R.C., D.H.B.; Visualization - R.S.; Supervision - M.R.C., D.H.B.; Project Administration - M.R.C., D.H.B. ; Funding acquisition - R.S., A.M.N., M.R.C., D.H.B.

## Extended Data

**Figure 1-1.**
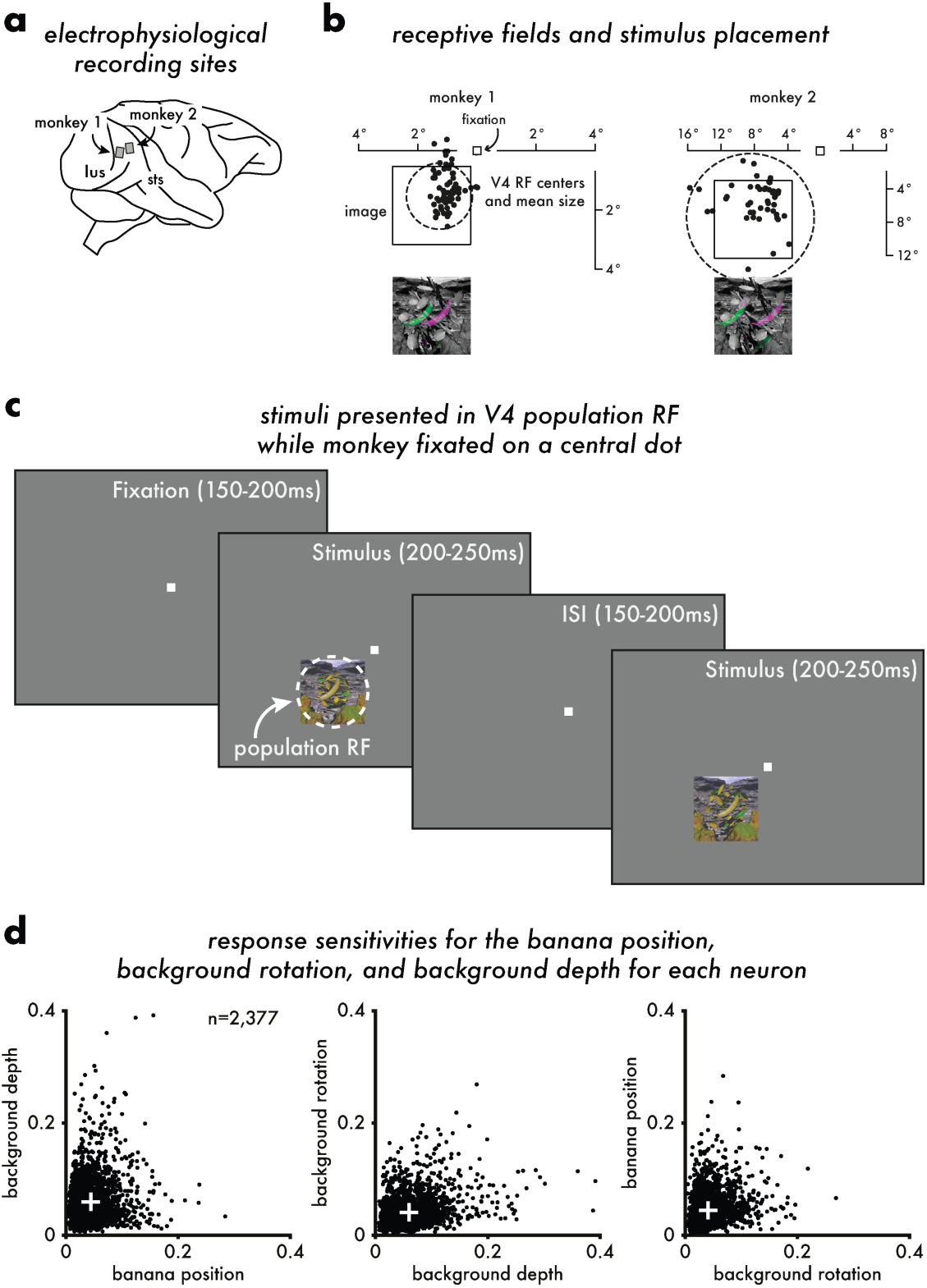
Recording locations, stimulus positions, behavioral task, and unit responsivity for monkeys. **a:** Multielectrode arrays (96 channels each) were chronically implanted in V4 of two monkeys. **b:** The receptive fields of the recorded multiunits were mapped using Gabor and two-dimensional shape stimuli. The estimated receptive field (RF) center of each visually responsive multiunit is depicted by a black point. The mean size of the RF for each population is depicted by the dashed circle. The black square indicates the position of the image on the screen. The size and position of the images for each session were chosen such that all the variations in stimulus position and background depth were within the estimated RF of the recorded V4 population. The variation in object position is indicated by the false color image in each panel. **c:** While monkeys fixated a central dot, stimuli were flashed on (200-250 ms) and off (150-200 ms) up to eight times before the monkey received a juice reward. Each of the 125 images was repeated between 8-10 times in every session. Electrophysiological recordings were collected using multielectrode arrays implanted in V4. Receptive fields (RFs) were mapped in an independent experiment using Gabor and 2D shape stimuli. Images were placed such that the variation in the central object (the banana) and background overlapped a large majority of RFs. **d:** Comparison of response sensitivity for banana position, background depth, and background rotation. Here, sensitivity is defined as the modulation index *(_rmax_-r_min_)/(r_max_+r_min_)* where *r_max_* and *r_min_* are respectively the maximum and minimum responses to variation in the corresponding parameter. The white cross represents the mean of the sensitivities across all visually responsive multiunits (n=2377) across all sessions.

**Figure 2-1:**
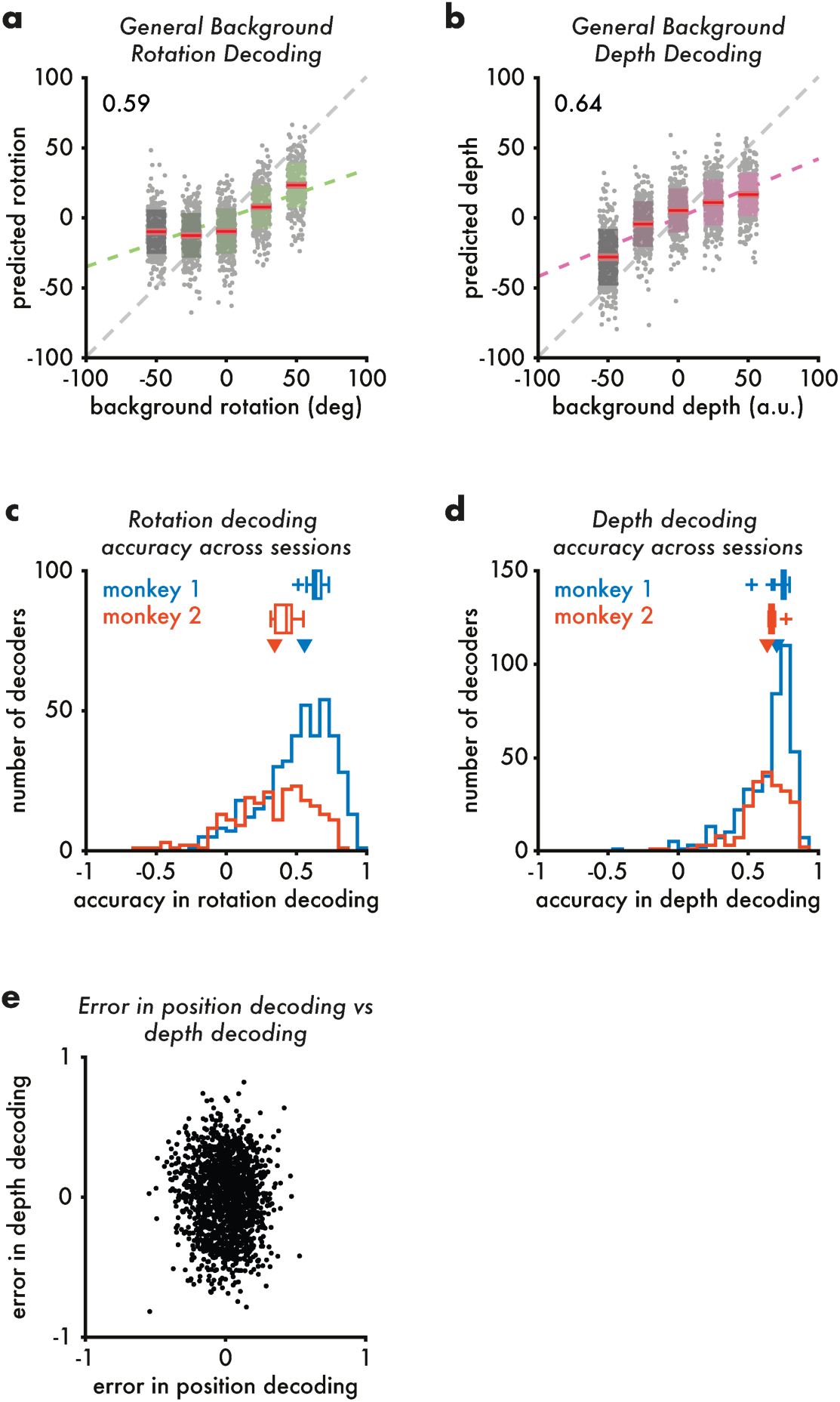
Background depth and rotation can also be decoded well from V4 population responses. Compare with Figure 2. **a:** Same as Figure 2b but for background rotation. Here, a general linear decoder was used to estimate the background rotation value in the face of variation in background depth and banana position. **b:** Same as Figure 2b but for background depth. Here, a general linear decoder was used to estimate the background depth value was decoded in the face of variation in background rotation and object position. **c:** Same as Figure 2c for specific background rotation decoding (each session contributes 25 values to the distribution). Here, the background rotation value was decoded while ignoring the other parameter variations across images. Arrows represent median decoding accuracy (0.558 for monkey 1, 0.344 for monkey 2). Box plots above the histograms show distribution of general decoder performance. **d:** Same as Figure 2c for specific background depth decoding (each session contributes 25 values to the distribution). Arrows represent median decoding accuracy (0.703 for monkey 1, 0.634 for monkey 2). Box plots above the histograms show distribution of general decoder performance. **e:** Comparison of errors for general decoding of object position and general decoding of background depth across trials. Compare with Figure 2d.

**Figure 4-1:**
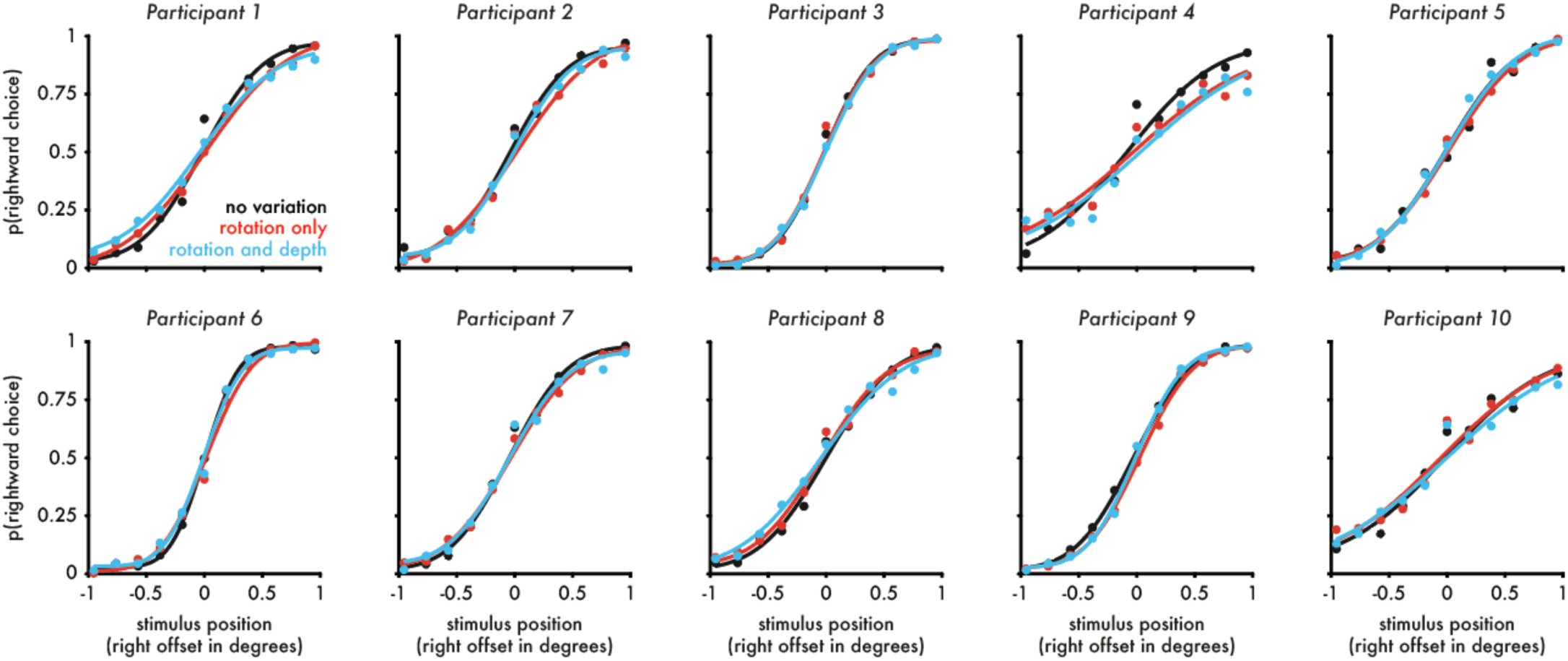
Psychometric functions for object position discrimination for each participant.

